# Identification of domestic cat hepadnavirus from a cat blood sample in Japan

**DOI:** 10.1101/2022.01.09.475482

**Authors:** Kazuki Takahashi, Yasuyuki Kaneko, Akiko Shibanai, Shushi Yamamoto, Ayana Katagiri, Tatsuyuki Osuga, Yoshiyuki Inoue, Kohei Kuroda, Mika Tanabe, Tamaki Okabayashi, Kiyokazu Naganobu, Isao Minobe, Akatsuki Saito

**Author notes:** Addresses correspondence to Yasuyuki Kaneko or Akatsuki Saito.

## Abstract

The hepatitis B virus (*Hepadnaviridae*) induces chronic hepatitis and hepatic cancer in humans. A novel domestic cat hepadnavirus (DCH) was recently identified in several countries, however, the DCH infection status of cats in Japan is unknown. Therefore, we investigated the DCH infection rate of 139 cat samples collected in Japan. We identified one positive blood sample (0.78%) from a 17-year-old female cat with chronically elevated alanine aminotransferase. Phylogenetic analysis demonstrated that the DCH strain identified in this study is genetically distinct from strains in other countries. Further investigations are required to elucidate the evolution of DCH and the impact of DCH infection on hepatic diseases in domestic cats.

## Main text

Hepatitis B virus (HBV) (*Hepadnaviridae*) is one of the causative agents of chronic hepatitis and hepatocellular carcinoma. Despite effective vaccines against HBV, over 350 million people worldwide are chronically infected with HBV [1]. One step in the life cycle of HBV involves the conversion of viral pregenomic RNA to a relaxed circular DNA; therefore, several reverse transcriptase inhibitors in combination with interferons are currently used for controlling HBV infection [2].

It has been unclear whether other mammals can be infected by similar hepadnaviruses that induce chronic hepatitis and hepatocellular carcinoma. In 2018, an Australian group identified a novel feline hepadnavirus, the domestic cat hepadnavirus (DCH), and since then, it has been detected in cats in several countries including Australia (6.5% of tested samples) [3] and Italy (10.8% of tested samples) [4]. Although it remains unclear whether DCH can induce hepatic diseases in cats, the detection of DCH has been associated with chronic hepatitis and hepatocellular carcinoma [5]. In addition, cats infected with the feline immunodeficiency virus (FIV) also tend to be positive for DCH [3], suggesting that immunodeficiency induced by FIV may lead to infection with DCH.

The fact that DCH has been discovered in several countries suggests that it is prevalent worldwide. Phylogenetic analyses in previous studies suggest that DCH is divergent among strains from different countries, suggesting that DCH has been evolving in each country. So far, DCH has been identified Australia [3], Thailand [6], Italy [5], the United Kingdom [7], and Malaysia [8]. Its presence and prevalence in Japan, however, remains undetermined. In this study, therefore, we investigated the presence and prevalence of DCH in Japan.

We screened 139 blood samples collected from several clinics in Japan. The blood samples were collected from cats that were mainly housed indoors. We used samples that were rest of blood testing for diagnosis. Prior to testing for DCH, we obtained the owner’s informed consent. This study was approved by the Animal Experiment Committee of the University of Miyazaki (authorization number: 2021-019). All experiments were performed in accordance with relevant guidelines and regulations.

All collected blood samples were mixed with the anticoagulants, heparin or ethylenedinitrilo-tetraacetic acid disodium (EDTA/2Na). Previous studies have demonstrated that heparin inhibits PCR [9], thus we used a KOD One PCR Master Mix (TOYOBO, Osaka, Japan), which is relatively tolerant to heparin contamination. To check the quality of the PCR, we used primer pairs to amplify the cat glyceraldehyde-3-phosphate dehydrogenase (*Gapdh*) housekeeping gene. The primers used to amplify the DCH genome are DCH-F: (5’-ATTCAAGCGCTCTATGAAGAGG-3’) and DCH-R: (5’-AAAAGTGAGGCAAGAGAGATGG-3’), while those used to amplify cat *Gapdh* are GAPDH-F: (5’-CCTTCATTGACCTCAACTACAT-3’) and GAPDH-R: (5’-CCCCAGTAGACTCCACAACATAC-3’). The PCR conditions were 40 cycles of 98°C for 10 sec, 60°C for 5 sec, and 68°C for 10 sec, followed by 68°C for 7 min. We used a synthesized DNA fragment encoding the partial DCH genome (Eurofins) as a positive control for DCH. The amplicons were visualized on a 1.5% agarose gel.

We detected *Gapdh* amplicons from the 129 samples tested. A single sample (#116) (0.78% of tested samples) was positive for DCH (**Figure 1**). We then sequenced the entire viral genome using six pairs of primers whose sequences are listed in **Table 1**. We designed the primers on the Primer3 website (https://bioinfo.ut.ee/primer3-0.4.0/). All six fragments were efficiently amplified. Each amplicon was run on an agarose gel and then extracted from the gel with QIAquick Gel Extraction Kit (Qiagen, Tokyo, Japan). The sequences of the amplicons were determined using a BigDye Terminator v3.1 Cycle Sequencing Kit (Thermo Fisher Scientific) on an Applied Biosystems 3130xl DNA Analyzer (Thermo Fisher Scientific). The sequence assembly was performed with 4Peaks (Nucleobytes) and Microsoft Word 2019 (Microsoft). The assembled sequence was deposited in GenBank (Accession# LC668427).

**TABLE 1.**
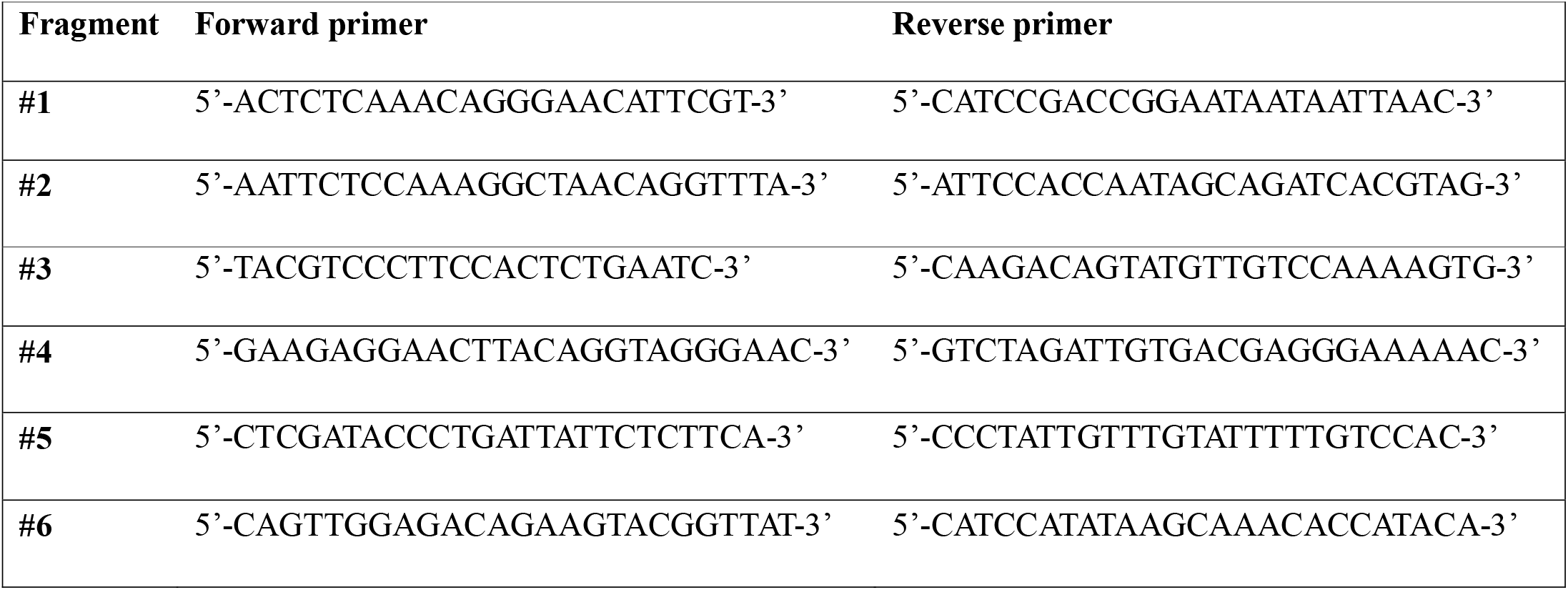
Primers used for determining viral whole genome.

**Fig. 1.**
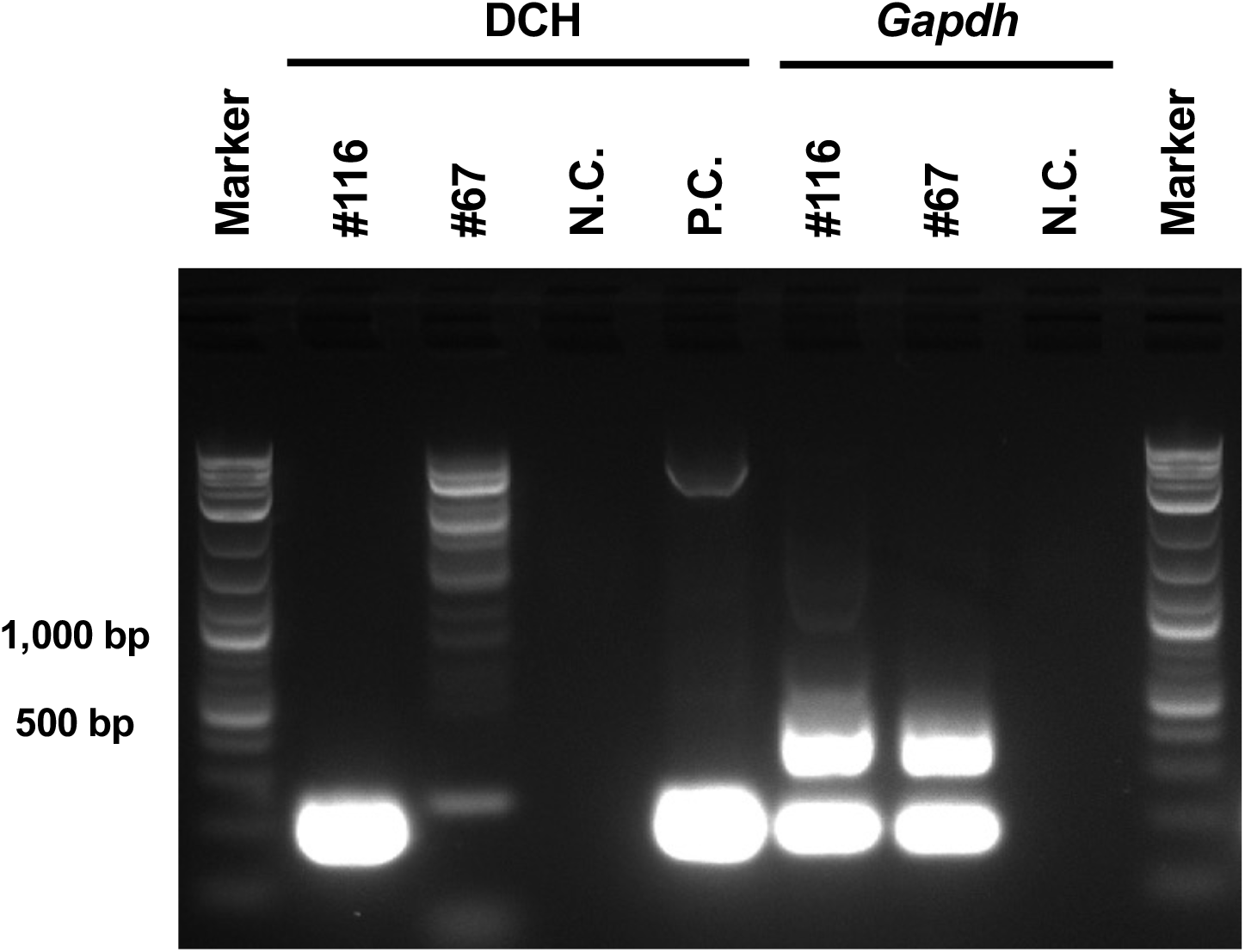
PCR screening of blood samples from cats to detect the domestic cat hepadnavirus (DCH) genome. Heparinized blood was used to amplify DCH and the cat *Gapdh* gene. Marker, N.C., and P.C. denote DNA size marker, negative control, and positive control, respectively.

The sequence of our strain was aligned with those of 15 other DCH strains and the sequences were analyzed phylogenetically using MEGA X (MEGA Software). Our phylogenetic analysis revealed that the amino acid sequences of the polymerase, surface, and core proteins of Domestic cat hepadnavirus Japan/KT116/2021 were genetically close to those of previously reported strains (**Figure 2**). In contrast, the amino acid sequence of the X protein of Domestic cat hepadnavirus Japan/KT116/2021 was distinct from those of other strains. The X protein of HBV (HBx) plays roles in silencing host antiviral defenses and promoting viral transcription [10, 11]. Therefore, it will be important to elucidate the impact of polymorphism in the DCH X protein.

**Fig. 2.**
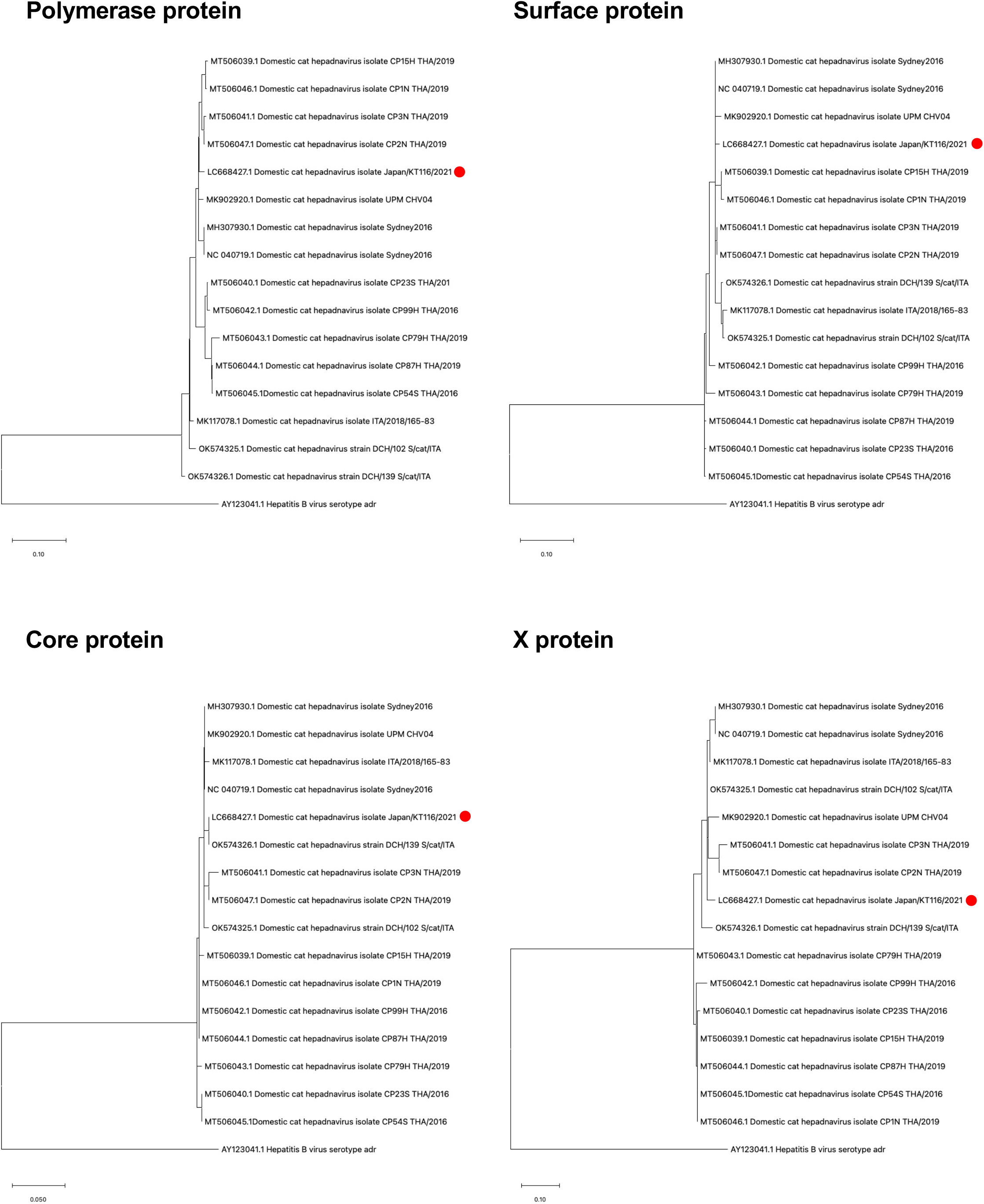
Phylogenetic analysis of DCH/Japan/KT116/2021. The phylogenetic position of DCH Malaysia within the family *Hepadnaviridae*. The maximum likelihood tree is based on the each hepadnaviral protein sequences retrieved from the GenBank database. The Domestic cat hepadnavirus Japan/KT116/2021 (Accession# LC668427) is indicated by a red dot. The tree is drawn to scale, thus the branch lengths correspond to the number of substitutions per site.

The source of the DCH-positive sample (#116) was a 17-year-old female cat (Cat #116) born in Japan and with no record of traveling overseas. Therefore, Cat #116 was likely infected with DCH in Japan. At the time of the PCR test, Cat #116 had no health problems, based on interviews and physical examinations. However, prior to its death due to acute neuropathy, we observed a persistent elevation of alanine aminotransferase (ALT) (**Figure 3**). Moreover, Cat #116 underwent a splenectomy one year before the PCR test, and was then diagnosed with a mast cell tumor. After this diagnosis, the cat was treated with CCNU, an anticancer drug. While we cannot exclude the possibility that the treatments against the mast cell tumor induced elevated levels of ALT, it is also possible that DCH infection had affected the health status of Cat #116. Currently, it is not known whether the mast cell tumor itself or the treatments against mast cell tumor had induced immunosuppression in Cat #116, leading to infection with DCH. As Cat #116 was tested positive for DCH after its death, we were unable to perform an autopsy. Note that Cat #116 was negative for FIV and feline leukemia virus (FeLV).

**Fig. 3.**
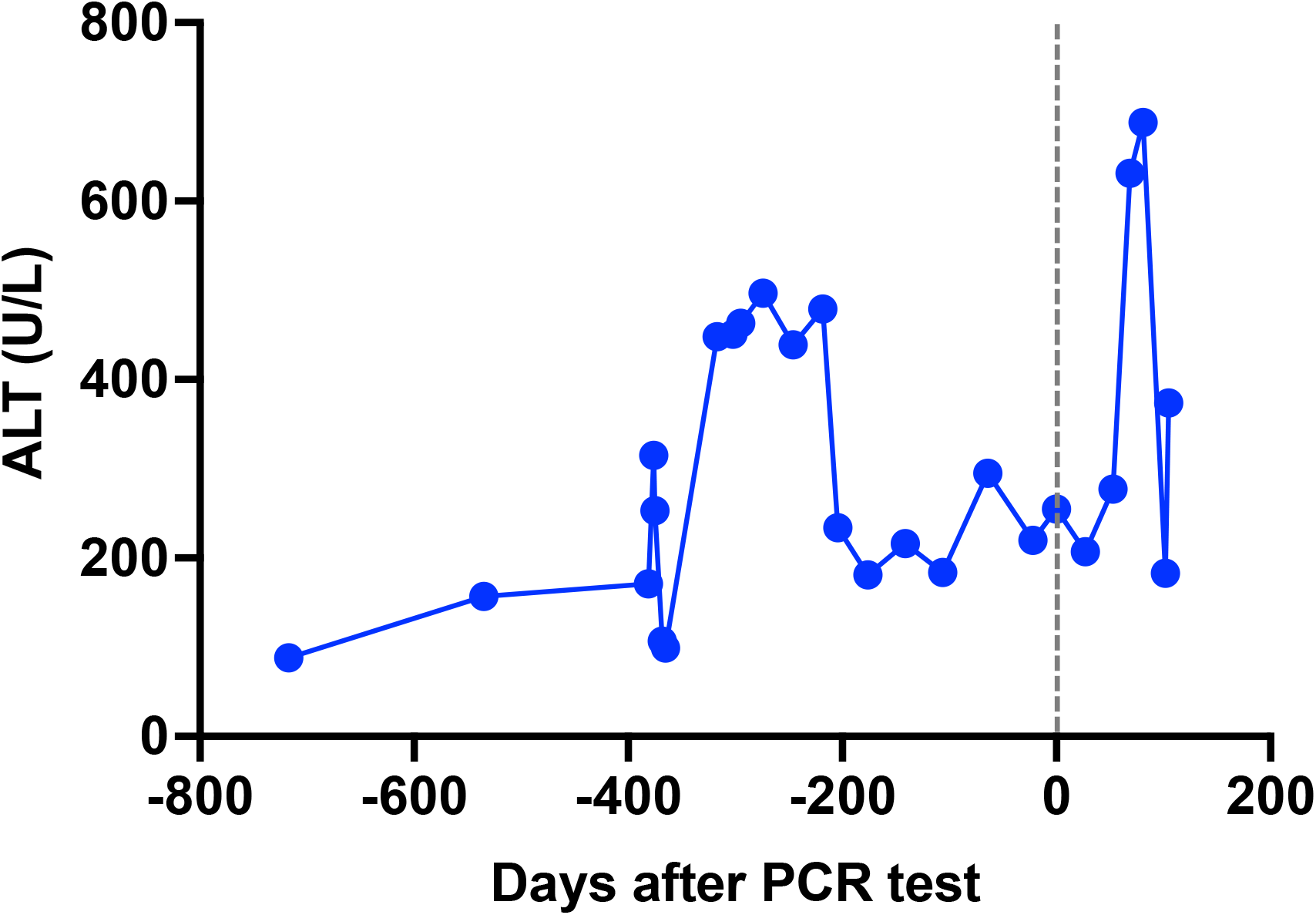
Changes in alanine aminotransferase (ALT) levels in Cat #116. A plot of ALT levels in each blood test. The X-axis denotes “Days after PCR test.”

In conclusion, to the best of our knowledge, this is the first time that DCH has been detected in a cat in Japan. The prevalence of DCH in this study (0.78%) is lower than those of previous studies in other countries [3, 4]. Further investigation is required to determine the reason for this difference. The results of phylogenetic analysis of the X protein reveal that Domestic cat hepadnavirus Japan/KT116/2021 is genetically distinct from previously described strains from other countries, suggesting that Domestic cat hepadnavirus Japan/KT116/2021 is a strain native to Japan. Because homologous recombination has been reported with HBV [12, 13] or DCH [6], it is important to monitor the infection status and evolutionary history of DCH in every country with a large domestic cat population. Also, the impact of DCH infection on chronic hepatitis in cats should be elucidated.

## ACKNOWLEDGMENTS

This work was supported by grants from Japan Society for the Promotion of Science (JSPS) KAKENHI Grant-in-Aid for Scientific Research (C) 19K06382 (to AS); KAKENHI Grant-in-Aid for Scientific Research (B) 21H02361 (to TO and AS); and from a Grant for Joint Research Projects of the Research Institute for Microbial Diseases, Osaka University (to AS). We thank Tomoko Nishiuchi for her support.

